# SynVerse: A Framework for Systematic Evaluation of Deep Learning Based Drug Synergy Prediction Models

**DOI:** 10.1101/2025.04.30.651516

**Authors:** Nure Tasnina, Maryam Haghani, T. M. Murali

**Affiliations:** Dept. of Computer Science, Virginia Tech, Blacksburg, VA 24060, USA

## Abstract

Synergistic drug combinations are often used to treat cancer. Experimental exploration of all possibilities is expensive. Deep learning (DL) for predicting the synergy of drug pairs in specific cell lines might provide an alternative. However, current approaches often suffer from data leakage. They also lack systematic ablation studies. To address these gaps, we propose SynVerse, a comprehensive evaluation framework featuring four data-splitting strategies to assess DL model generalizability and three ablation studies: module-based, feature shuffling, and a novel network-based approach to disentangle factors influencing model performance. We evaluated sixteen models incorporating eight drug- and cell line-specific features, five preprocessing techniques, and two widely used encoders. Our analysis revealed several insights. None of the models outperformed a naive baseline using one-hot encoding as features. Biologically meaningful drug or cell line features and drug–drug interactions were not the drivers of predictive performance. All models demonstrated poor generalization to unseen drugs and cell lines. SynVerse emphasizes the need for substantial improvements before computational predictors can reliably support experimental and clinical settings.

## 1 Introduction

Combination therapy is a cornerstone of cancer treatment, offering increased efficacy [1], reduced side effects [2], and the potential to overcome drug resistance compared to monotherapy [3, 4]. For instance, Sabutoclax, a small-molecule BH3 mimetic administered with Minocycline, a synthetic tetracycline, displays anti-tumour activity by reducing tumor growth *in vitro* and *in vivo* on pancreatic ductal adenocarcinoma [5]. This type of therapy relies on the “synergy” of drug combinations, which is the phenomenon where the combined effect of two or more drugs exceeds the expected additive effect [6, 7].

The synergy of a drug combination is usually first established in cancer cell lines during preclinical studies [8, 9]. The size of the drug space and variations in synergy from one cell line to another preclude the possibility of systematic experimental exploration of drug combinations. In recent years, many deep learning (DL)-based regression models have appeared to predict the synergy score for a given drug-drug-cell line triplet. The availability of open-access datasets of synergy scores such as O’Neil *et al*. [2], DrugComb [10], and Drug-CombDB [11] has facilitated the development of these methods. These databases provide different scoring methods for assessing the synergy of a drug combination, e.g., Loewe [12] and *S*_*mean*_ [13].

These DL models often use the encoder-decoder architecture, which consists of three principal modules: (i) Preprocessing: This optional module performs a one-time transformation on the raw features of drugs and cell lines; (ii) Learnable encoder: This optional module consists of a DL architecture that takes the preprocessed features of drugs and cell lines as inputs and generates feature representations, i.e., embeddings for drugs and cell lines; and Learnable decoder: This module consists of a multi-layer perception (MLP) that predicts the synergy score using the embeddings of drugs and cell lines as inputs. The encoder and decoder modules are trained in an end-to-end manner, minimizing the discrepancy between predicted and true synergy.

DeepSynergy [14] was the first DL model for synergy prediction, using chemical features of drugs and gene expression values in cell lines with an MLP-based decoder. It outperformed other baseline machine learning models, such as support vector machines, random forests, and XGBoost. Subsequent models have explored diverse encoder architectures including MLPs [15, 16], autoencoders [17, 18], graph neural networks [19, 18], and transformers [20, 21, 22, 18, 23]. These models leverage a range of feature types, including chemical fingerprints [15, 17, 16], SMILES (i.e., text-based notation for representing chemical structures) [19, 21, 22, 23], gene expression and multi-omic data [17, 20, 18, 16, 23], and drug target profiles [20]. Most models are trained and evaluated on the O’Neil *et al*. dataset, with some using DrugComb [15, 16, 23].

Despite the publication of over 20 DL-based synergy prediction methods, several key issues remain. A crucial aspect is the assessment of model generalizability, i.e., its performance on unseen drug pairs, cell lines, and drugs. Unfortunately, many studies [24, 25, 26, 27, 28], including an evaluation framework for DL-based models [29], have primarily split triplets randomly between training and test sets. This approach can lead to data leakage due to the presence of the same drug pairs in both the training and test sets [30]. Additionally, studies often lack thorough ablation analysis to investigate the contributions of individual drug and cell line features [14, 15, 21, 19] or the individual components of models [20, 19]. This omission is particularly concerning since recent advances in architecture and feature selection have increased model complexity but only modestly improved performance over DeepSynergy. For instance, TranSynergy [20] and SynergyX [23] reported improvements as small as 1% and 2%, respectively, in terms of Pearson’s correlation coefficient (PCC) (Figure 1A). Adding to this concern are the inconsistent performance of DeepSynergy across studies (Figure 1A) and the variations in synergy score distributions (Figure 1B), even when methods use the same datasets and use identical synergy measures, e.g., MARSY [16] and SynergyX [23]. These gaps make it difficult to assess whether a model’s performance reflects genuine improvements in feature and model selection or is simply an artifact of the dataset used and preprocessing, highlighting the need for a rigorous, first-principles examination of the most common components of DL-based synergy prediction methods.

**Figure 1.**
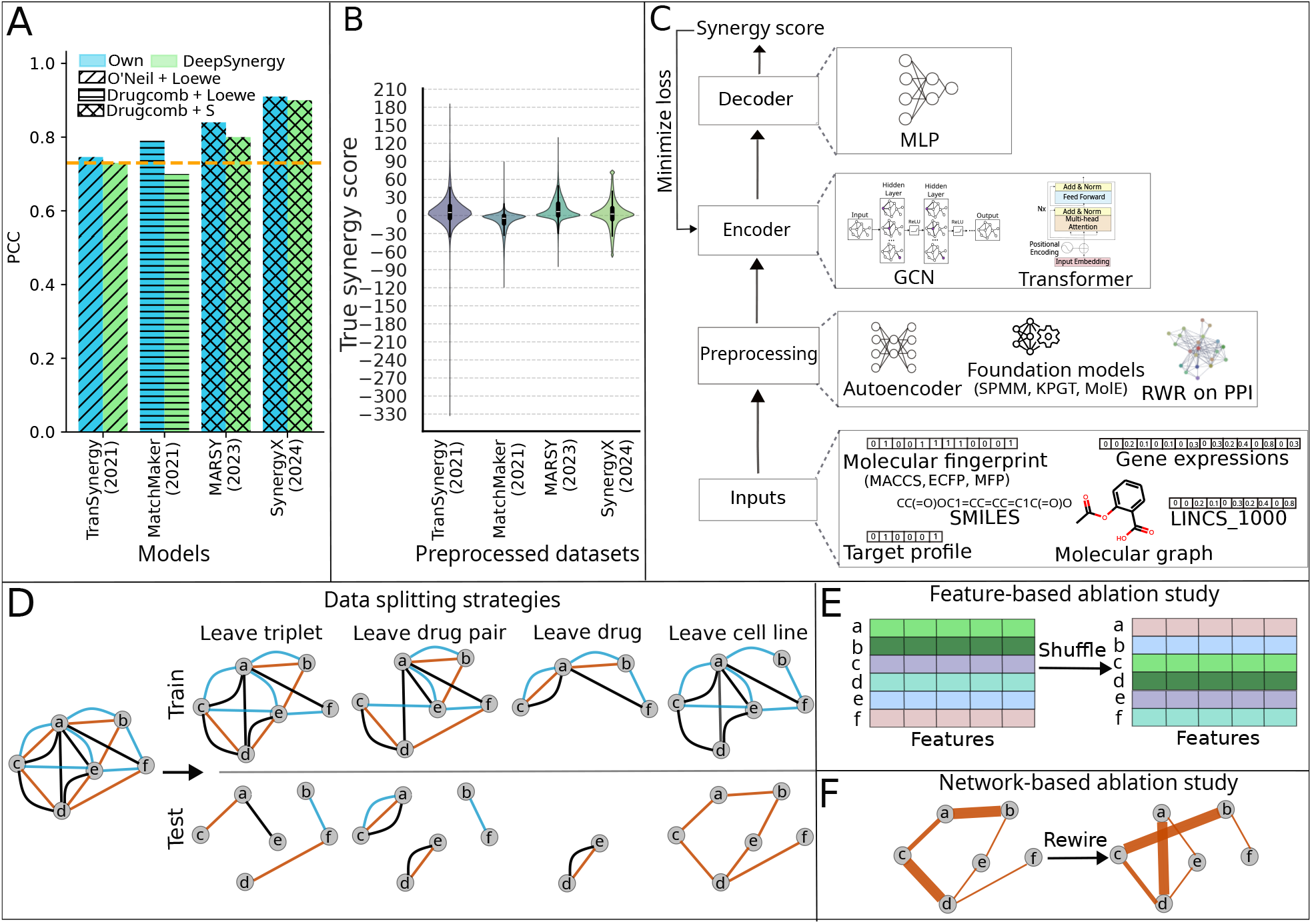
SynVerse: Motivation, framework, and evaluation. **A.** Comparison of existing models with DeepSynergy in predicting synergy of triplets with unseen drug pairs. Each pair of bars (in two colors) represents the following: 1) the model’s own performance (blue) 2) DeepSynergy’s performance reported by the publication for the model (green). The orange dotted line represents the performance of DeepSynergy reported in the original publication [14]. The different textures (/, -, x) of the bar plots indicate the combination of the dataset and synergy score considered by the corresponding models. **B**. Distribution of synergy scores. The *x*-axis represents the models and the *y*-axis the synergy score distribution in the dataset preprocessed by a model. **C**. SynVerse architecture composed of preprocessor, encoder, and decoder. The left panel illustrates the generic architecture of a model provided in SynVerse. The right panel showcases the the options provided in each module. **D**. Four data splitting strategies incorporated in SynVerse. We represent the triplets with synergy scores as a network. Each node in the network corresponds to a unique drug. An edge between two drugs represents the synergy between them in a particular cell line, indicated by the color of the edge. **E**. A pictorial depiction of shuffling used in our feature based ablation study. **F**. A pictorial depiction of network rewiring that preserves node strength (weighted degree) used in our network based ablation study. The thickness of an edge is directly proportional to its synergy score.

Inspired by these observations, we propose an extensive evaluation strategy for DL-based synergy prediction models, enabling a systematic assessment of model generalizability while supporting rigorous ablation studies. Our strategy includes four data-splitting methods: (1) leave triplet, (2) leave drug pair, (3) leave drug, and (4) leave cell line, which evaluate how well a model generalizes to unseen triplets, drug pairs, drugs, and cell lines, respectively (Figure 1D). These splitting strategies enable the evaluation of models under controlled data leakage conditions. We incorporate three types of ablation studies to evaluate the contributions of different components of the model architectures and inputs. First, we include a conventional module-based ablation analysis to assess the impact of individual modules on overall performance. Second, we introduce a shuffling-based feature ablation approach, where each drug (or cell line) is assigned the representation of a randomly selected counterpart (Figure 1E). Third, we design a novel network-based ablation strategy. Synergy score-labeled triplets form a weighted network, where nodes represent drugs and edge weights denote synergy scores. This ablation assesses how much predictive power arises from the topology of the network as opposed to the input features themselves (Figure 1F).

To embody this evaluation strategy, we design SynVerse, a framework with an encoder-decoder architecture (Figure 1C). SynVerse incorporates diverse input features and a reasonable approximation of model architectures commonly employed by current DL-based synergy prediction methods. Rather than assessing pre-existing synergy prediction architectures, we developed SynVerse to maximize flexibility in feature selection and model design. Two key considerations motivated this decision. First, directly integrating existing models into the evaluation would inherently limit the ability to explore different feature-model combinations because of the constraints imposed by specific implementations. Second, our literature review showed that even the most recent models often fail to outperform DeepSynergy, a simple MLP-based synergy prediction model [20, 23]. This lack of improvement underscores the importance of systematically investigating various feature representations and machine learning architectures, an effort facilitated by SynVerse.

Our systematic evaluation of sixteen synergy prediction models encompassed eight drugs and cell line features, five preprocessing techniques, and two DL-based encoders (Figure 1C). Our findings were unexpected and illuminating. When tested on triplets containing unseen drug pairs, all the models achieved good performance in terms of Pearson’s correlation coefficient (between 0.78 and 0.94). However, none of them outperformed a naive *baseline* model that employed one-hot encoding for both drugs and cell lines and a simple MLP-based architecture. Next, models with shuffled drug and cell line features performed comparably to those using the original representations, suggesting that model performance was not primarily driven by biologically, chemically, or cell line-specific informative features. Moreover, our network-based ablation study indicated that the models predominantly learned patterns in the synergy score distribution of *individual* drugs, while largely disregarding the partner drug in the triplet. This finding suggests that the models were not leveraging meaningful network topological information from the drug-drug synergy network to drive their performance. Finally, we assessed the predictive performance of the models in triplets containing previously unseen drugs and cell lines. All models exhibited poor performance in this setting, highlighting their lack of generalizability.

## 2 Results

### 2.1 Overview of Experimental Setup

#### Features

We incorporated three different molecular descriptors for drugs, i.e., Molecular Access System (MACCS), Extended Connectivity Fingerprint (ECFP), and Morgan fingerprints (MFP). We also used molecular structure, SMILES, and drug targets (Section 4.1) as drug features. As cell line features, we included the gene expression profiles of untreated cell lines (Genex) (Section 4.1). Additionally, we filtered out the gene expression data to contain expression for 978 landmark genes mentioned in the LINCS L1000 project [31] (LINCS 1000) (Section 4.1). The terms inside the parentheses represent the names we use for these features in the rest of the paper.

#### Model architectures

We implemented three preprocessing techniques: 1) Auto-encoder-based dimensionality reduction (AE), applied to sparse, binary features such as MACCS, ECFP, Morgan fingerprints, and one-hot encoding; 2) Random walk with restart (RWR) on protein-protein interaction networks, applied to binary drug target profile; and 3) Molecular foundation models, where we generated embeddings from SMILES representations using three pretrained molecular foundation models, i.e., MolE [32], KPGT [33], SPMM [34]. We integrated two DL-based encoders to learn embeddings for drugs: i) a graph convolutional neural network (GCN) [35] to generate embeddings from the molecular graph of a drug, and ii) a transformer, which encoded the text-based SMILES representation of a drug.

#### Evaluation metrics and datasets

We evaluated model performance using two standard metrics for regression tasks: (1) Root Mean Squared Error (RMSE) and (2) Pearson’s Correlation Coefficient (PCC). For each architecture, we reported the performance of the best model, selected through hyperparameter tuning (Section 4.7).

We trained and evaluated the models to predict the *S*_*mean*_ synergy score for drug–cell line triplets extracted from DrugComb [10] (Section 4.2, Supplementary Note 1.1). We trained and evaluated each model only on the subset of triplets for which the required features were available for both drugs and the corresponding cell line.

### 2.2 Evaluation Results and Analysis

SynVerse offers flexibility in selecting any combination of the drug and cell line features present in the framework. In this work, we focus on evaluating models that use one type of (drug or cell line) feature at a time (Section 4.6). We adopt this approach to specifically assess the utility of individual features in predicting synergy. We denote each model in the format *x* (*p, q*) where *x* is the feature, *p* is the preprocessor, *q* is the encoder (Table 1) and *p* and *q* are optional. For instance, a model denoted as *SMILES (Transformer)* employs SMILES as the drug feature with a Transformer-based encoder and one-hot encoding (default) for cell line features. As the GCN and Transformer-based encoders are designed for graph and text-formatted inputs, respectively, and our integrated preprocessing methods yield numerical vectors, our analysis considered models that use either a preprocessing method or an encoder for a certain feature, but not both.

**Table 1.** Specification of models. The combination of features, preprocessing techniques, and encoders evaluated by SynVerse.

| Model name | Drug feature |  |  | Cell line feature |  |  |
| --- | --- | --- | --- | --- | --- | --- |
|  | Feature name | Preprocessing | Encoder | Feature name | Preprocessing | Encoder |
| MACCS | MACCS | - | - | One hot | - | - |
| MACCS (AE) | MACCS | Autoencoder, Standardization | - | One hot | Autoencoder, Standardization | - |
| ECFP | ECFP_4 | - | - | One hot | - | - |
| ECFP (AE) | ECFP_4 | Autoencoder, Standardization | - | One hot | Autoencoder, Standardization | - |
| MFP | Morgan Fingerprint | - | - | One hot | - | - |
| MFP (AE) | Morgan Fingerprint | Autoencoder, Standardization | - | One hot | Autoencoder, Standardization | - |
| Mol Graph (GCN) | Molecular structure | - | GCN | One hot | - | - |
| SMILES (Transformer) | SMILES | - | Transformer | One hot | - | - |
| SMILES (SPMM) | SMILES | SPMM | - | One hot | - | - |
| SMILES (KPGT) | SMILES | KPGT | - | One hot | - | - |
| SMILES (MolE) | SMILES | MolE | - | One hot | - | - |
| Target | Drug target | - | - | One hot | - | - |
| Target (RWR) | Drug target | RWR on PPI network | - | One hot | - | - |
| Target (AE) | Drug target | Autoencoder, Standardization | - | One hot | Autoencoder, Standardization | - |
| Genex | One hot | - | - | Gene expression | Standardization | - |
| LINCS_1000 | One hot | - | - | Expression of 978 Landmark genes | Standardization | - |
| Baseline | One hot | - | - | One hot | - | - |

First, we present each model’s performance in predicting *S*_*mean*_ synergy score for triplets with unseen drug pairs. Next, we compare the results to the baseline model (Figure 2A). These two analyses comprise the first ablation study, where we seek to determine the contribution of each component of the architecture. We then present the feature-based ablation study (Figure 2B) and the findings from network-based ablation (Figure 2C). Finally, we explore results from two more data-splitting strategies, i.e., leave drug and leave cell line (Figure 3). We present the results using the leave triplet strategy in Supplementary Note 1.2, as this split allows the highest overlap between training and test sets, leading to a higher risk of data leakage.

**Figure 2.**
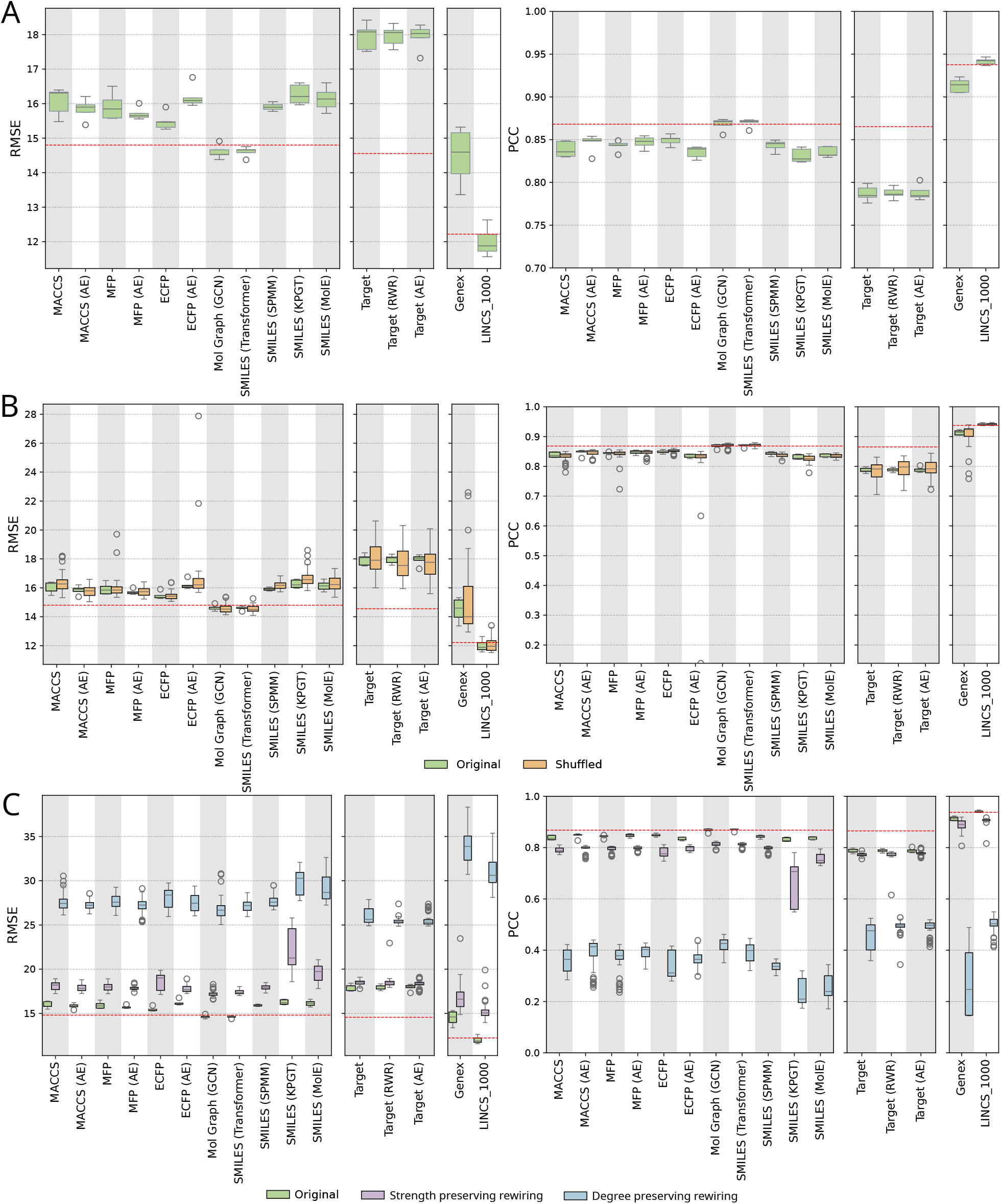
Evaluation of models on test data using leave drug pair split. **A**. Distribution of RMSE (left panel) and PCC (right) between true and predicted scores, across five independent runs. The *x*-axis represents the models and the *y*-axis the RMSE or PCC. The red dotted line indicates the median performance of the baseline model. **B**. Performance comparison between models with original vs. shuffled features. **C**. Performance comparison between models trained on original vs. rewired network.

We organize each set of results in three subplots, each containing a group of models categorized by the features they employ. The first group includes models that use SMILES or SMILES-derived features such as molecular fingerprints and molecular structures. The second group consists of models that use drug targets as input features, and the third group includes models that rely on gene expression data from cell lines. This grouping ensured that all models within a group were trained and evaluated on the same subset of triplets.

#### Performance of models on unseen drug pairs

We aimed to evaluate the utility of individual drug and cell line features in predicting synergy on triplets with unseen drug pairs, using a leave drug pair split (Figure 2).

First, we considered the 11 models that used SMILES or SMILES-derived features (Figure 2A). They showed a significant difference in their performance (*p*-value of 4.99 × 10^−5^ for RMSE, *p*-value of 4.88 × 10^−5^ for PCC; Kruskal-Wallis test). Among them, *Mol Graph (GCN)* and *SMILES (Transformer)* achieved the lowest median RMSE (14.52 and 14.65, respectively) and the highest median PCC (0.87 for both). Both of them performed significantly better than the other models (*p*-value *<* 0.03 for both RMSE and PCC; one-sided Mann-Whitney U test with Benjamini-Hochberg correction). There was no significant difference between these two models (*p*-value of *>* 0.8 for both RMSE and PCC; two-sided Mann-Whitney U test with Benjamini-Hochberg correction). Note that *Mol Graph (GCN)* generated drug embeddings from molecular graphs using a GCN, while *SMILES (Transformer)* derived embeddings from SMILES using a transformer-based encoder.

Next, we examined the models exploiting drug target information in different forms, i.e., binary target profile, target profile generated by running RWR on PPI network, and autoencoder (Figure 2A). These model were similar to each other in performance with median RMSE ranging from 18.03 to 18.08 and PCC of 0.79 (*p*-value *>* 0.93 for RMSE and PCC; Kruskal-Wallis test).

Finally, among the models utilizing gene expression data, *LINCS 1000*, which uses expression from 978 landmark genes, achieved the best performance with a median RMSE of 11.88 and a PCC of 0.94 (Figure 2A). It significantly outperformed the *Genex* model, which leverages expression data from all available genes (median RMSE of 14.6, PCC of 0.91; *p*-value = 0.004; one-sided Mann-Whitney U test with Benjamini-Hochberg correction).

Across all the 16 models, the lowest PCC was 0.79. Overall, we concluded the models appeared to exhibit moderate to strong performance in predicting synergy for unseen drug combinations.

#### Comparison to baseline

To investigate whether the model’s predictive power stems from the model architecture and features used, we compared each model to a baseline that employed one-hot encoding for drug and cell line features and an MLP as the predictor. Notably, this baseline did not incorporate any biologically meaningful features. All models, except *Mol Graph (GCN), SMILES (Transformer)*, and *LINCS 1000*, performed worse than the baseline (Figure 2A). Even for these three models, the improvement over the baseline was not statistically significant (adjusted *p*-value = 1.0; one-sided Mann-Whitney U test with Benjamini-Hochberg correction).

#### Feature-based ablation study

The comparable or superior performance of the baseline raised concerns about whether the models relied on biologically meaningful drug and cell line features. To investigate this phenomenon further, we shuffled drug (and cell line) features, assigning each drug (and cell line) the features of another at random. For each original feature set, we generated 10 shuffled versions to account for variability and repeated model training and evaluation for each. We observed no significant difference between the performance of models trained on shuffled features and those trained on the original features (Figure 2B), with *p*-values ranging from 0.49 to 0.94 for RMSE and 0.85 to 0.97 for PCC (two-sided Mann-Whitney U test with Benjamini-Hochberg correction). This observation reinforced the conclusion that the observed performance of the models might not be driven by biologically meaningful features.

#### Network-based ablation study

Since the models did not perform better than baselines or alternatives with shuffled features, we sought to understand the basis for their observed performance (Figure 2A). We considered triplets (with synergy scores) from each cell line as a weighted network, where each node represents a drug and each weighted edge corresponds to the synergy score of the adjacent drugs. To assess whether and to what extent models utilized this network topology, we conducted a systematic network-based ablation study. First, we rewired each cell line-specific training network using a strength (i.e., weighted degree) preserving method (Section 4.5), maintaining the total sum of positive and negative synergy scores for each drug while randomizing drug-drug pairs. Next, we rewired the networks preserving only degree sequences instead of strength (Section 4.5). We then trained models on these randomized triplets and evaluated them on the original test data. We randomized each training network 10 times using each rewiring method.

Models trained on the original networks consistently outperformed those trained on either type of rewired networks, with *p*-value *<* 0.05 for both RMSE and PCC (one-sided Mann-Whitney U test with Benjamini-Hochberg correction) (Figure 2C). Notably, models trained on randomized, strength-preserved networks retained between 85% and 99% of the performance (in terms of PCC) observed with the original networks. This result suggests that models may primarily rely on the distribution of strength or synergy scores associated with individual drugs but disregard the partner drugs in the triplets. The substantially lower performance observed for models trained on degree-preserved randomized networks, which retained only 25% to 63% of the original performance, further supports this interpretation. These findings are concerning as they imply that the models are not effectively capturing biologically meaningful drug interactions to inform their predictions.

#### Performance of models on leave drug split

With the aim of assessing the models’ capability to generalize towards unseen drugs, we trained and evaluated models using the leave drug strategy (Figure 3A). The models leveraging molecular fingerprints, SMILES representations, or molecular graph exhibited poor performance (median RMSE between 27.13 and 34.3 and median PCC between 0.1 and 0.21) in predicting synergy for triplets containing unseen drugs. The models utilizing drug targets features also performed poorly with the lowest median RMSE of 30.25 achieved by *Target (RWR)*. The highest PCC of 0.14 was achieved by autoencoder based model *Target (AE)*. Moreover, when compared to the baseline, none of these models showed significant improvement (*p*-value *>* 0.43; one-sided Mann-Whitney U test with Benjamini-Hochberg correction. Notably, in this split, the performance of each model exhibited substantial variability across runs, as reflected by the wider interquartile range. These findings indicate that the models struggled to capture generalizable patterns from the chemical features of drugs. Consequently, when faced with unseen drugs, the models performed poorly compared to their performance on the leave drug pair split (Figure 2A), where all drugs in the test set were also present during training.

**Figure 3.**
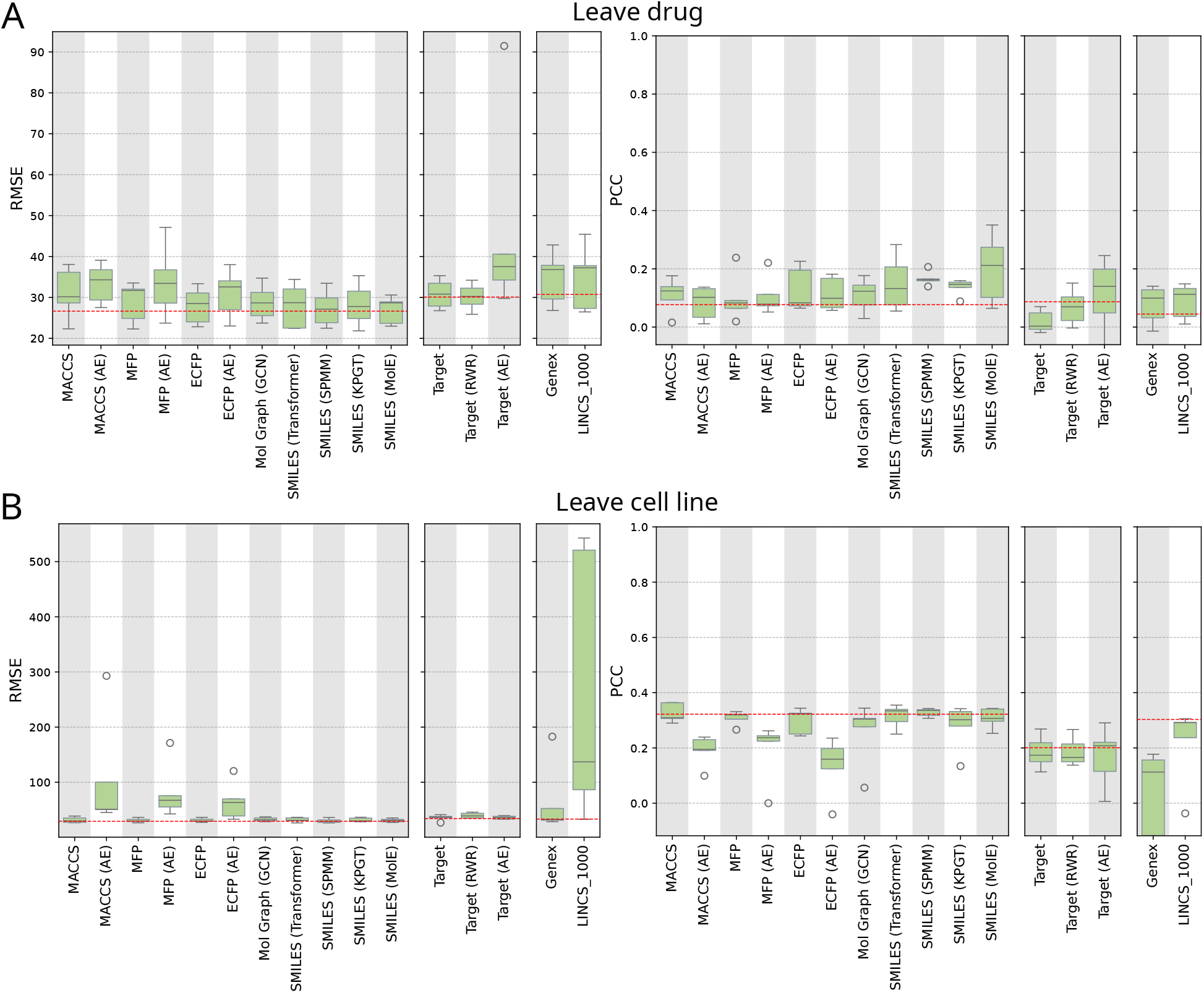
Evaluation of model generalizability on unseen drugs and cell lines. **A**. Performance of models on test dataset based on leave drug split. **B**. Performance of models on test dataset based on leave cell line split.

#### Performance of models on leave cell line split

We evaluated each model’s ability to predict synergy scores for triplets containing unseen cell lines (Figure 3B). All models exhibited poor performance. *A priori*, we expected that models with gene expression-based features may generalize well to unseen cell lines in the test dataset. However, their performance fell short, with a median RMSE of 32.33 and PCC of 0.11 for *Genex* and RMSE of 136.94 and PCC of 0.29 for *LINCS 1000*. Moreover, when compared to the baseline, none of these models showed significant improvement(*p*-value = 1.0; one-sided Mann-Whitney U test with Benjamini-Hochberg correction). These findings indicate that the models struggled to derive generalizable patterns from the cell line features. Consequently, when faced with novel cell lines during testing, the models performed poorly.

## 3 Discussion

In this study, we present a systematic evaluation strategy for DL-based synergy prediction models, addressing critical gaps in the literature related to data leakage and ablation studies. To assess model performance under controlled data leakage conditions, we incorporated four data splitting strategies designed to evaluate performance on previously unseen triplets, drug pairs, individual drugs, and cell lines. We also devised three ablation studies to disentangle the contributions of model architecture, feature representation, and network topology. We implemented SynVerse, a comprehensive framework that assessed sixteen models incorporating eight drug-and cell line-specific features, five preprocessing techniques, and two widely used encoders in synergy prediction research.

At first glance, all models appeared to exhibit moderate to strong performance, with PCC ranging from 0.78 to 0.94 when tested on predicting *S*_*mean*_ score for triplets with unseen drug pairs. However, none of these models significantly outperformed a baseline model that used one-hot encodings of features, i.e., it did not include any biologically meaningful features. To further contextualize the performance of our baseline, we compared it to MARSY and SynergyX, two state-of-the-art models (Supplementary Note 1.3). To this end, we trained and evaluated our baseline on the synergy datasets processed in the corresponding publications. Remarkably, the baseline closely matched the performance of MARSY and SynergyX, with PCC values differing by at most 0.03.

Further investigation through feature-based ablation studies confirmed that the models in the SynVerse framework failed to capture meaningful semantic patterns from drug and cell line features, as shuffling these features had no detrimental effect on performance. Additionally, network-based ablation experiments suggested that the models were leveraging topological shortcuts, specifically, the strength distribution of the nodes (drugs) in the input network instead of biologically meaningful drug-drug interactions to drive their predictions. Consequently, when tasked with predicting synergy for triplets containing novel drugs and cell lines, model performance dropped drastically, underscoring their inability to extract biologically relevant patterns from input features.

These findings align with recent studies on link prediction in protein-protein interaction networks [36] and biological knowledge graphs [37] in which improper partitioning of training and test data has been shown to artificially inflate model performance. Furthermore, Chatterjee *et al*. ([38]) showed that multiple state-of-the-art models for protein-ligand binding prediction maintained their performance even when input features were randomized. Through further analysis, they revealed that these models primarily exploited a topological shortcut, specifically, the degree information of proteins and ligands within the drug–target interaction network. This observation aligns with the findings of Geirhos *et al*. [39], who reported that deep learning models frequently rely on shortcuts present in training data to achieve high predictive performance.

Our systematic evaluation strategy mitigates the risk of reporting inflated model performance due to data leakage and provides insights into the factors driving model performance. Our findings underscore that computational predictors must be substantially improved before their results achieve a quality sufficient to enable their translation to experimental and clinical settings. In the future, we aim to extend our framework to evaluate multitask synergy prediction models, where models are trained not only to predict drug synergy but also to perform related tasks, such as estimating individual drug sensitivity scores [40, 16]. A systematic evaluation of these multitasking models could provide valuable insights into their predictive performance.

## 4 Materials and Methods

In this section, we first describe the datasets we use to create drug and cell line features. Then we present the dataset for synergy scores. Finally, we explain the architecture of SynVerse and our proposed evaluation strategy.

### 4.1 Feature Extraction

The features we use fall into three broad categories: i) SMILES and SMILES-derived features such as drug molecular fingerprints and molecular graph; ii) drug targets; and iii) gene expression-based features for cell lines. We now describe how we process each feature in detail.

#### Drug identifier

This is a one-hot encoding-based binary feature vector of length equal to the number of drugs considered in SynVerse.

#### SMILES

SMILES (Simplified Molecular Input Line Entry System) [41] is a compact line notation system that represents the structure of chemical compounds using short ASCII strings. It encodes molecular structures, including atoms, bonds, and connectivity. We obtained the SMILES representation for each drug by mapping its name to a PubChem compound using the PubChemPy package [42, 43].

#### Molecular descriptors

These features capture various structural and chemical properties of molecules. Among these, two-dimensional (2D) molecular descriptors represent structural information derived from the 2D representation of molecules, such as connectivity indices and structural fragments. Two commonly used types of 2D molecular descriptors are structural keys and hashed fingerprints. Structural keys encode the structure of a molecule into a binary bit string, where each bit represents a predefined structural feature (e.g., substructures or fragments such as a C=N group or a six-membered ring). If a molecule contains a specific feature, the corresponding bit is set to 1; otherwise, it is set to 0. However, structural keys are limited to capturing only those features that are predefined in the fragment library. In contrast, hashed fingerprints do not rely on predefined fragment libraries. Instead, they are generated by systematically enumerating all possible fragments of a molecule up to a specified size and then converting these fragments into numeric values using a hashing function. This method allows hashed fingerprints to encode a broader range of molecular features, providing greater flexibility.

In this study, we utilized three molecular descriptors, encompassing both structural keys and hashed fingerprints, which have been employed in existing drug synergy prediction models [14, 17]: 1) Molecular Access System (MACCS): A structural key represented as a binary vector of length 166, encoding predefined molecular features [44]. 2) Morgan fingerprint: A hashed fingerprint with a radius of 2 and a dimension of 256 [45]. 3) Extended Connectivity Fingerprint (*ECFP*_4_): A hashed fingerprint with a radius of 2 and dimension 1, 024 [46]. We computed all these descriptors from the corresponding SMILES representations of molecules using RDKit [47].

#### Drug targets

We downloaded information on drug targets from the Therapeutic Target Database (TTD) [48] where drug targets are curated through a systematic process involving the collection of newly approved drugs, clinical trial drugs, preclinical drugs, and patented drugs from various authoritative sources, including ClinicalTrials.gov, company reports, patents, and literature. Each drug’s therapeutic target is validated based on its functional role in disease phenotypes and its ability to achieve therapeutic efficacy. As of the 2024 update, TTD includes 3, 730 targets and 39, 862 drugs. However, only 1, 425 drugs present in the preprocessed synergy dataset (see Section 4.2 below) had their target information available in TTD. These drugs had 2, 536 unique targets in TTD. Hence, for each drug, we computed a target profile of dimension 2, 536, a binary vector where each dimension had the value one if and only if the drug acts upon the corresponding target.

#### Molecular graph

We used RDKit [47] and DeepChem [49] to convert the SMILES representation of a drug into a graph where each node is an atom and each edge is a bond between two atoms.

#### Cell line identifier

This feature is a one-hot encoding, where each index represents a cell line. For a specific cell line, only its corresponding index is 1, while the rest are 0.

#### Gene expression values in untreated cell lines

We utilized cell line-specific messenger RNA expression data obtained from the Cell Line Encyclopedia (CCLE)[50]. Of the cell lines included in the preprocessed synergy dataset, expression data were available for only 136 cell lines, encompassing a total of 20, 068 genes. From this data, we constructed two types of input features: (i) the full gene expression profile, using all available genes, and (ii) a reduced-dimensional representation based on a subset of biologically informative genes.

For the second type, we followed the approach adopted in previous synergy prediction models [15, 21]. We used the gene expression data only for 978 landmark genes mentioned in the LINCS L1000 project [31]. These landmark genes have been shown to capture 82% of the information present in the full transcriptome [51]. Among these, 963 genes overlapped with those available in the CCLE expression dataset.

#### Protein-protein interaction network

We used a human protein-protein interaction network from STRING (version 12) ([52]) comprising 12, 650 nodes and 197, 784 edges applying the “highest” score cutoff of 900 on the confidence score of edges. We describe how we compute features from this network when we present the pre-processing module of SynVerse.

### 4.2 Synergy Dataset Preparation

DrugComb is one of the largest publicly available datasets providing synergy scores for drug-drug-cell line triplets calculated using various metrics. These synergy measures are generally based on the deviation of the observed response from a theoretical model. In this study, we focused on *S*_*mean*_ [13], a recently-developed measure that evaluates synergy by analyzing the deviation of the drug combination sensitivity score (CSS) from a reference model. This reference model predicts the expected percentage inhibition effect using monotherapy doseresponse data. The CSS is computed from dose-response curves of drug pairs tested in a cross-design setup, where each drug is combined at varying doses with a fixed dose of a background drug. This methodology enables the concurrent assessment of both sensitivity and synergy, facilitating the identification of efficacious and synergistic drug combinations for cancer treatment. Moreover, drug combinations deemed synergistic by *S*_*mean*_, which employs IC50 concentrations for the background drugs, are therapeutically more relevant since such combinations avoid higher concentrations, often associated with undesirable off-target effects and side effects [13].

The DrugComb dataset was not readily usable for model training due to drug name inconsistencies and variability among triplet replicates. To address these challenges, we implemented a series of preprocessing steps. First, we removed samples containing responses to single drugs, retaining a total of 751, 498 drug combinations derived from 4, 269 unique drugs tested across 295 cell lines. To ensure consistency in drug identification, we standardized drug names by mapping them to PubChem Compound Identifiers (CIDs), a non-zero integer that uniquely identifies a chemical structure in PubChem [43]. During this process, we excluded triplets involving compounds without assigned CIDs (86 drugs) or those with multiple CIDs linked to differing SMILES representations (can be indicative of different salt forms) (87 drugs). To address inconsistencies in synergy scores across replicates for a triplet, we imposed a threshold on the standard deviation of scores. We excluded triplets with a standard deviation exceeding 0.1; only 7% of the triplets failed to meet this threshold. We calculated a consensus synergy score for each remaining triplet by averaging the scores of its replicates.

After these steps, the dataset comprised 556, 905 triplets involving 3, 969 unique drugs across 263 cell lines. Since the features used varied from one model to another, we constructed three subsets of triplets from this dataset, each corresponding to one of the three feature categories. The first subset included only those triplets for which SMILES strings (or features derived from them) were available for both drugs. The second subset comprised triplets with drug target information. The third subset contained triplets with gene expression data available for the corresponding cell lines. Next, to ensure that each cell line was adequately represented during training, we required that each cell line appear in at least 5% of the triplets in its respective subset. Finally, SMILES-based subset contained 105, 722 triplets, 2, 582 drugs, and 9 cell lines; the target-based subset included 53, 850 triplets, 275 drugs, and 13 cell lines; and the gene expression-based subset comprised 104, 093 triplets, 194 drugs, and 15 cell lines (Supplementary Note 1.1).

### 4.3 SynVerse Framework

We formulate synergy prediction as a regression problem where the goal is to predict the synergy score for a triplet (*d*_*i*_, *d*_*j*_, *c*_*k*_) utilizing the features of the drugs *d*_*i*_ and *d*_*j*_ and the cell line *c*_*k*_. As an in-depth literature review revealed a common encoder-decoder-based architecture used by DL-based synergy prediction models, our framework SynVerse consists of three principal modules: (i) **A preprocessing module** that takes raw features of drugs and cell lines as inputs; (ii) **A learnable encoder module** that takes preprocessed features of drugs and cell lines as inputs and generates embedding for drugs and cell lines; and (iii) **A learnable decoder module** that predicts the synergy score given embeddings for a pair of drugs and a cell line as inputs (Figure 1). We train the encoder-decoder modules in an end-to-end manner, minimizing the discrepancy between predicted and true synergy.

#### 4.3.1 Preprocessing Module

The preprocessing module allows for additional processing after feature extraction. It is not involved in the end-to-end training. We provide the following options for preprocessing in SynVerse:

##### Autoencoder

Many of the extracted features, such as drug target profiles and molecular fingerprints, are sparse binary vectors. Hence, we provide a preprocessing step involving an autoencoder to generate a low-dimensional, dense feature representation. After training, we extracted the embedding from the bottleneck layer (the layer with the lowest dimension) of the autoencoder.

##### RWR on PPI

Given the proteins targeted by a drug and a protein-protein interaction network, we compute a numerical drug target profile using the random walk with restart (RWR) [53] algorithm. By treating the drug targets as restart nodes, we calculated the probabilities of a random walker reaching each node in the network at steady state. We used the corresponding vector of node probabilities as the profile of the drug. This approach is motivated by the need to capture the drug’s impact on non-target proteins.

##### Pretrained molecular foundation models

Recent advancements in molecular foundation models, pretrained on extensive molecular datasets, have demonstrated their capability to learn robust, generalizable, and informative representations of drugs. We used three such pretrained foundation models to investigate whether their embeddings can enhance synergy prediction, : KPGT [33], SPMM [34], and MolE [32].

Knowledge-guided Pre-training of Graph Transformer (KPGT) [33] is a self-supervised learning framework to provide generalizable and robust molecular representations. KPGT combines the Line Graph Transformer (LiGhT), tailored for molecular graph structures, with a knowledge-guided pre-training strategy to capture both structural and semantic knowledge. Pre-trained on a large dataset of around 2 million molecules, KPGT demonstrated superior performance in molecular property prediction and practical applicability in drug discovery. Building on the strengths of multimodal learning, Structure-Property Multi-Modal foundation model (SPMM) [34] integrates molecular structures and biochemical properties using a Transformer architecture. SPMM extracts intramodal features and performs intermodal fusion via self-attention and cross-attention mechanisms, respectively. SPMM is pretrained on 50 million SMILES representations from PubChem [54] and 53 molecular properties calculated with RDKit [47]. SPMM has demonstrated its ability to generalize as a foundation model by performing well on bidirectional tasks such as property prediction (SMILES-to-properties) and property-conditioned molecule generation (properties-to-SMILES) and uni-modal tasks, including molecule classification and reaction prediction.

MolE (Molecular Embeddings) [32] is another foundation model with a transformer-based architecture. MolE is trained on molecular graphs from approximately 842 million molecules using a self-supervised pretraining strategy that enabled the model to learn by predicting the environment (i.e., the atom type and connectivity of all neighboring atoms) around each atom. It adapted the disentangled attention mechanism from DeBERTa ([55]) to incorporate relative positional information between atoms within a molecular graph. These innovations enabled MolE to produce embeddings that were well-suited for various downstream tasks. However, since the version of MolE pretrained on 842 million molecules was unavailable, we utilized a smaller model provided by the authors, which was pretrained on the GuacaMol dataset [56] comprising approximately 1.2 million compounds.

#### 4.3.2 Encoder Module

The Encoder module contains two options for learnable DL-based models.

##### Graph convolution neural network (GCN)

To compute feature representations that capture the structural information of a drug, we utilized a Graph Convolutional Neural Network (GCN) based encoder. The input to GCN is a molecular graph, where nodes represent the atoms in a drug molecule, edges represent the chemical bonds between atoms. In addition, the GCN takes as input a feature matrix containing the initial representation of the nodes. We used RDKit to convert the SMILES format of a molecule into a molecular graph. We employed DeepChem [49] to compute atom-level features for each drug.

In each layer of the GCN, the process of learning drug representation entails message passing between each node and its neighboring nodes, producing node-level feature vectors using the following equation.

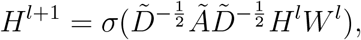

where *Ã* ∈ *R*^*n*×*n*^ is the adjacency matrix of the molecular graph containing *n* nodes and includes self-loops, 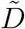 is a diagonal matrix with 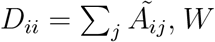, *W* is a learnable weight matrix, and *H*^*l*^ ∈ *R*^*n*×*c*^ is the learned feature matrix at layer *l*, and *σ* is an activation function (e.g., rectified linear unit (ReLU)). After the final GCN layer, we aggregated the learned atom-level features through a global Max Pooling layer to generate a graph-level feature vector representing the entire drug molecule.

##### Transformer

We used a Transformer-based encoder to learn long-range dependencies and contextual information in molecular structures represented by SMILES. First, we added a [CLS] token at the beginning and a [SEP] token at the end of each SMILES string. Then, we tokenized each SMILES string by mapping each subword to an integer ID, adapting the vocabulary provided in SPMM [34], which in turn derived from a pretraining SMILES corpus using the byte pair encoding (BPE) algorithm [57]. We then passed the integer tokens through an embedding layer, which mapped them into continuous embeddings represented as real-valued vectors. To encode positional information, we implemented two approaches: learnable encodings (via an embedding layer trained with the model) and fixed encodings (using a sinusoidal function). We treated the choice of type as a hyperparameter [58]. We then passed the input embeddings and positional encodings through a stack of Transformer encoder layers. The Transformer encoder transforms the initial token embeddings into more informative, context-aware representations, using the attention mechanism which captures contextual relationships by focusing on different parts of the sequence, and a feedforward neural network that introduces non-linearity [58, 59]. After the input embeddings are processed by the Transformer layers, the model generates an output embedding for each token. We used the processed embedding of the [CLS] token as the overall representation of the drug molecule.

#### 4.3.3 Decoder Module

We concatenated the learned embeddings of the drugs and the cell line, obtained from the encoder module, and used them as input to an MLP-based regression model to predict the synergy score. During training, we minimized the mean squared error (MSE) between the predicted and true synergy scores.

### 4.4 Data Splitting Strategies

To train and evaluate our models, we employed the following four data splitting strategies to ensure that the model is evaluated on its capability of predicting synergy for unseen triplets, drug pairs, drugs, and cell lines.

1. **Leave triplet**. We split the drug-drug-cell line triplets uniformly at random into train (80%) and test (20%) sets. This approach evaluates how a model performs when it sees a new triplet.
2. **Leave drug pair**. We considered the set of unique drug pairs (ignoring the cell lines in a triplet) and split them uniformly at random into two sets *T*_1_ (80% of drug pairs) and *T*_2_ (20% of drug pairs). We formed the training (respectively, test) set by including all triplets that involved a drug pair in *T*_1_ (respectively, *T*_2_). Hence, we excluded any drug pair present in the test set from the training set, i.e., if a triplet (*d*_*i*_, *d*_*j*_, *c*_*k*_) appeared in the test set, all triplets involving the drug pair (*d*_*i*_, *d*_*j*_) were also in the test set. Unlike the “leave triplet” split, this strategy ensures that no drug pair appears in both the training and test sets.
3. **Leave drug**. Analogous to “leave drug pair”, we partitioned the set of unique drugs into two sets *T*_1_ and *T*_2_. We formed the training (respectively, test) set by including all triplets where both drugs were present in *T*_1_ (respectively, *T*_2_). Consequently, no drug present in the test set appeared in the training set, i.e., if a triplet (*d*_*i*_, *d*_*j*_, *c*_*k*_) was present in the test set, then all triplets with either *d*_*i*_ or *d*_*j*_ were also in the test set. Note that this split excluded triplets in which one drug appeared in *T*_1_ and the other in *T*_2_. This strategy allows us to assess a model’s capability in predicting synergy when encountering a novel drug.
4. **Leave cell line**. Here, we split the triplet by cell lines. Hence, any cell line present in the test set was not present in the training set, i.e., if (*d*_*i*_, *d*_*j*_, *c*_*k*_) was present in the test set, then all triplets from cell line *c*_*k*_ were also in the test set. This strategy evaluates a model’s predictive performance on novel cell lines.

### 4.5 Ablation Studies

To investigate the factors that drive a synergy predictor’s performance, we performed the following three ablation studies.

1. **Comparison with baseline**: To evaluate the impact of the model’s features and its architecture, we designed a comparison with a baseline that employed one-hot encodings for drug and cell line features and an MLP as the decoder. A model with effective features and architecture should outperform this baseline that lacks biologically meaningful features and relies on a simple MLP.
2. **Feature shuffling:** We used this strategy to assess whether a model captures biologically meaningful patterns in drug and cell line features. Specifically, we randomly reassigned feature representations across entities, thereby disrupting any underlying biological associations. For instance, in the case of the *SMILES (Transformer)* model, we shuffled the SMILES strings among drugs, i.e., we assigned each drug the SMILES string for a different drug, uniformly selected at random. We trained and evaluated the model using these shuffled features. If the model’s performance remains unaffected by this shuffling, it suggests that the model does not rely on the biological relevance of the features. Instead, it may treat the feature values merely as identifiers, similar to one-hot encoding.
3. **Network rewiring:** To assess the extent to which a model relies on the network topology of drug-drug-cell line triplets, we conducted a network-based ablation study. This experiment stems from the hypothesis that if models exploit node degree (weighted or unweighted) as a predictive shortcut rather than learning meaningful semantic patterns, their performance should remain relatively stable when the network’s connections are randomized while preserving node degrees [37].

For each cell line, we constructed a separate network where each node represents a drug and an edge weight corresponds to the synergy score between the adjacent drugs. This edge weight can be positive or negative. We employed two types of rewiring strategies that preserved the original network properties to different extents while randomizing the edges.

First, we employed the Maslov–Sneppen rewiring method [60] that preserves the degree sequence, i.e., the number of edges connected to each node, without considering edge weights (Supplementary Note 1.4). Given an undirected graph *G* = (*V, E*) with node set *V* and edge set *E*, this method maintains the degree *d*(*v*) of each node *v* ∈ *V* by randomly swapping edge pairs while ensuring no self-loops or multi-edges are introduced.

Second, we applied a simulated annealing-based method [61], which preserves both the degree and the strength, i.e., the sum of adjacent edge weights, of each node while rewiring the edges (Supplementary Note 1.4). Given a weighted graph *G* = (*V, E, w*), where *w* : *E* → ℝ maps each edge to a real-valued weight, this algorithm seeks to preserve the strength *s*(*v*) = ∑_*u*∈*N*(*v*)_ *w*_*vu*_ for each node *v*, where *N* (*v*) denotes the set of neighbors of *v*. Using simulated annealing, this method minimizes the mean squared error between the strength sequences of the original and rewired networks.

In both cases, for each cell line-specific network, we randomized the positive and negative edges separately to preserve the original degree distributions or strength distribution of each signed subgraph. Note that this approach introduced the possibility that the same pair of drugs in a randomized network could be connected by two edges, one with positive and the other with negative weight. However, since our hypothesis posits that models relying on shortcuts learn from preserved degree or strength rather than the specific edges, the appearance of such edges should not substantially affect the validity of our conclusions.

### 4.6 Evaluated Models

A user of SynVerse has the flexibility to choose any combination of drug and/or cell line features along with compatible preprocessing techniques and encoders implemented in SynVerse. For our analysis, though, we focused on the subset of models that used only one feature, optionally applying a preprocessing step and a single encoder to generate the final embedding (Table 1).

### 4.7 Hyperparameter Tuning

We used BOHB [62], a robust hyperparameter optimization method, to navigate the extensive hyperparameter space of our models, which included architectural parameters (e.g., the number and width of layers), optimization parameters (e.g., learning rate), and regularization parameters (e.g., dropout rates) (Table 2). BOHB efficiently identifies high-performing configurations by combining the rapid exploration capabilities of HyperBand [63] with the convergence guarantees offered by Bayesian Optimization [64]. Instead of relying on random sampling, BOHB incorporates a Bayesian optimization component to guide the search effectively. For our experiments, we utilized the Python library HpBandSter, developed by the authors of BOHB.

**Table 2.**
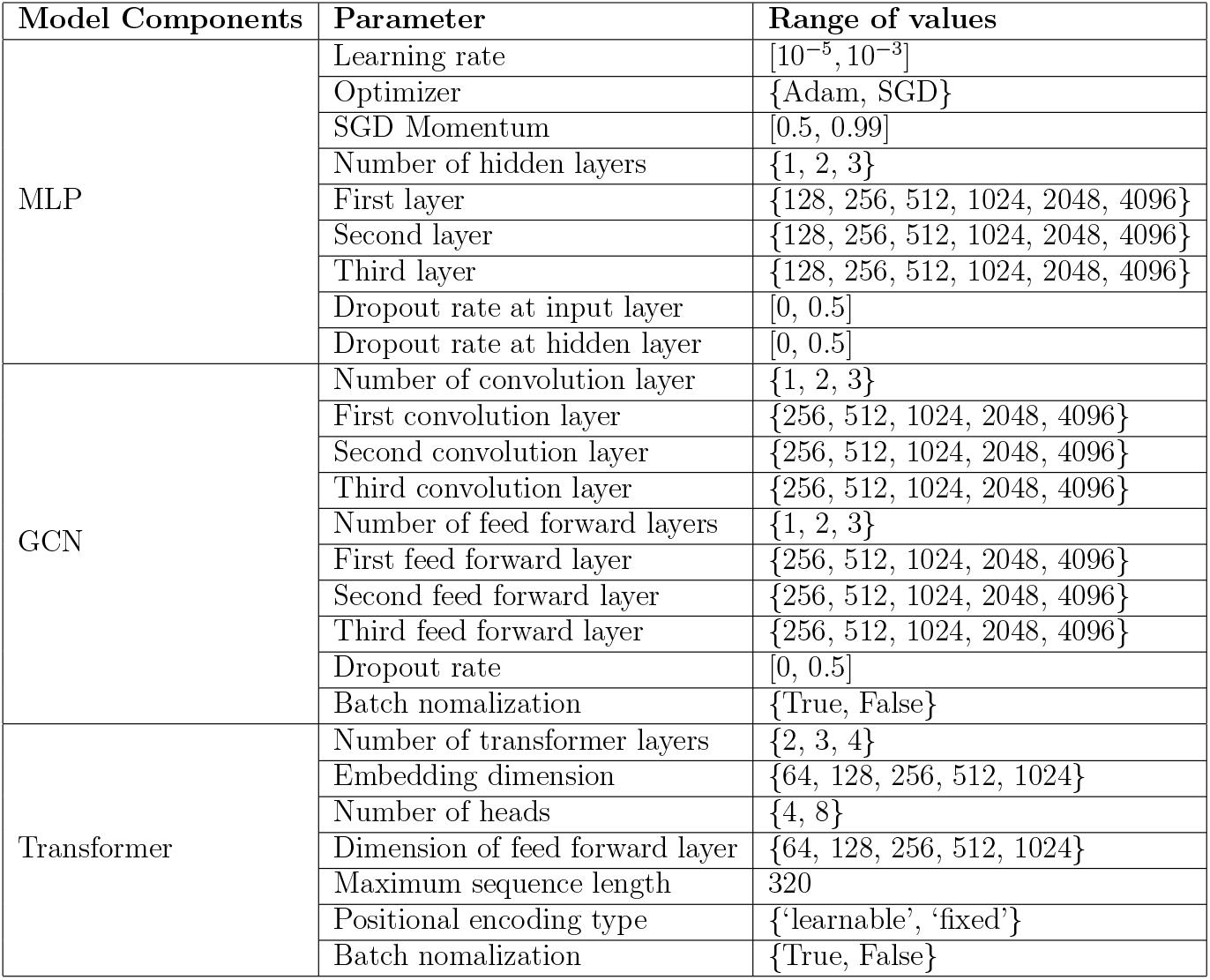
Hyperparameter configuration explored by BOHB for different models.

We performed hyperparameter tuning using a validation set comprising 25% of the training triplets. We applied early stopping with a patience of 90 epochs. We also used an adaptive learning rate strategy, ReduceLROnPlateau, which decreased the learning rate by a factor of 0.5 if the validation loss did not improve for 20 consecutive epochs.

## 5 Data Availability

All data used in this study are publicly available. The preprocessed data used in SynVerse are available on the Zenodo repository at https://zenodo.org/records/15277144.

## 6 Code Availability

The code to reproduce results, along with documentation and usage examples, is available on GitHub at https://github.com/Murali-group/SynVerse.

## 7 Acknowledgments

This work was supported by the National Science Foundation award DBI-2233967 and the College of Engineering at Virginia Tech.

