## Supplementary File for "SynVerse: A Framework for Systematic Evaluation of Deep Learning Based Drug Synergy Prediction Models"

### 1 Supplementary Notes

#### 1.1 Dataset Statistics: $S_{mean}$ Score

To facilitate the training and evaluation of models using different features, we constructed three subsets of triplets from DrugComb. The first subset included only those triplets for which SMILES strings (or features derived from them) were available for both drugs. The second subset comprised triplets with drug target information. The third subset contained triplets with gene expression data available for the corresponding cell lines. We imposed a minimum threshold of 5% on the abundance of triplets per cell line in the final dataset used for SynVerse, allowing different cell lines to qualify for inclusion across subsets.

SMILES-based subset contained $105,722$ triplets, $2,582$ drugs, and 9 cell lines; the target-based subset included $53,850$ triplets, 275 drugs, and 13 cell lines; and the gene expression-based subset comprised $104,093$ triplets, 194 drugs, and 15 cell lines. In the SMILES-based subset, the cell line *kbm7* was the most represented, appearing in $38,228$ triplets (38% of the subset), whereas *wm115* had the fewest, with $5,365$ triplets (5%) (Supplementary Figure 1A). The $S_{\text{mean}}$ score in this subset had a mean of 15.01 and a standard deviation of 29.27 (Supplementary Figure 1B).

Within the drug target-based subset, *kbm7* again had the largest presence, contributing $15,226$ triplets (28%), while *istmel1* was the least represented with $2,775$ triplets (5%) (Supplementary Figure 1A). This subset showed a similar distribution of synergy scores, with a mean of 15.76 and a standard deviation of 29.23 (Supplementary Figure 1B).

In contrast, the gene expression-based subset had a different distribution: *uacc257* contributed the highest number of triplets at $9,620$ (9%), and *colo792* had the lowest with $5,356$ (5%) (Supplementary Figure 1A). This group displayed a slightly higher average synergy score of 17.19, accompanied by a larger standard deviation of 36.25 (Supplementary Figure 1B).

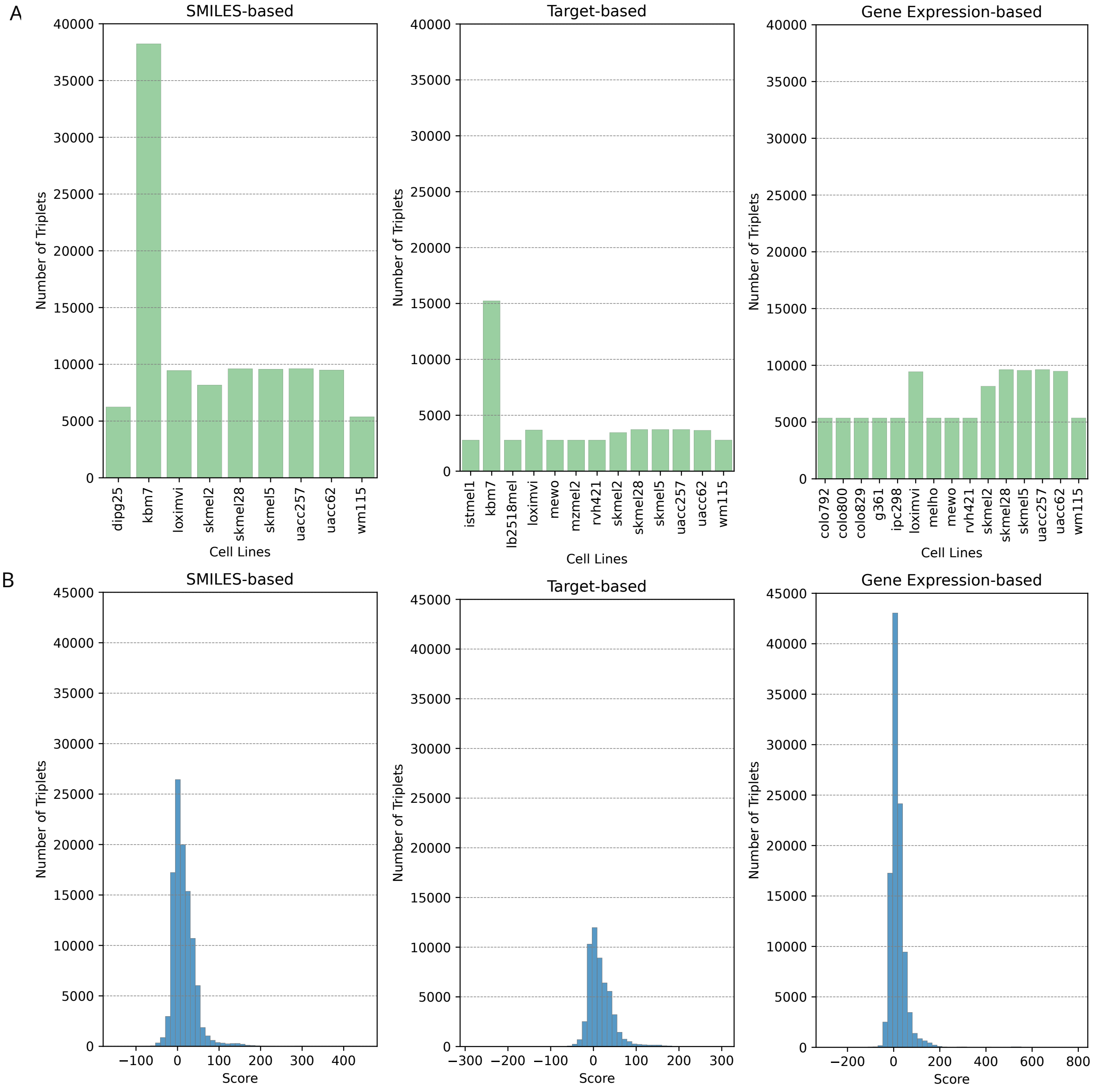


Supplementary Figure 1: **Distribution of $S_{mean}$ score in the DrugComb dataset**. **A.** The $x$-axis shows the names of cell lines, and the $y$-axis indicates the number of drug–drug–cell line triplets associated with each cell line. **B.** The $x$-axis displays binned $S_{mean}$ synergy scores, and the $y$-axis shows the number of triplets falling within each score bin.

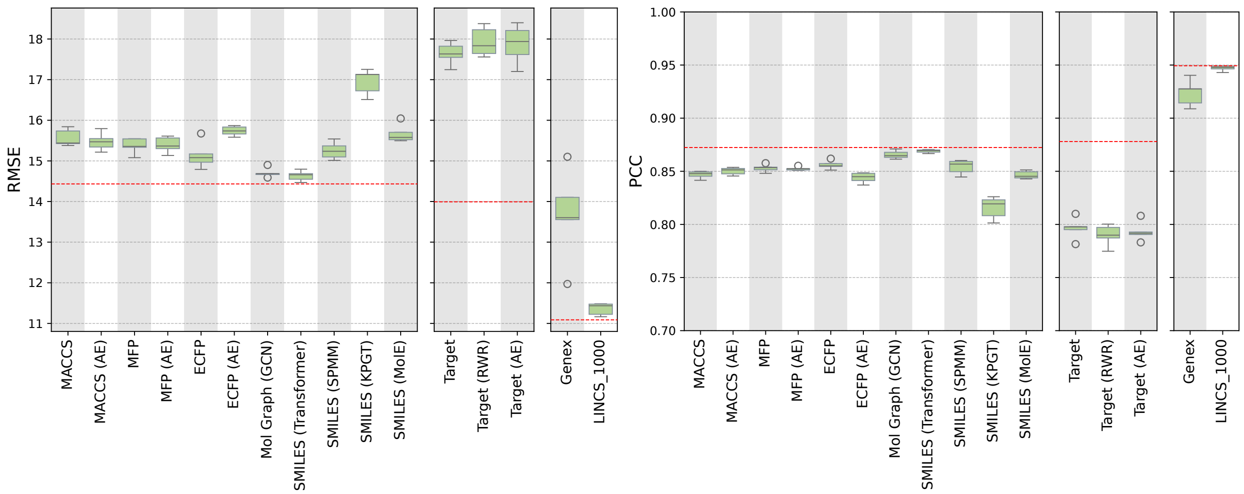


Supplementary Figure 2: **Evaluation of models in predicting $S_{mean}$ score using leave triplet split.** The $x$-axis represents the models, the $y$-axis displays RMSE (left panel) and PCC (right) between true and predicted scores by each model across five independent runs. The red dotted line indicates the median performance of the baseline model.

#### 1.2 Performance of Models Predicting $S_{mean}$ Using Leave Triplets Strategy

We aimed to assess the predictive value of individual drug and cell line features for synergy prediction on unseen drug–drug–cell line triplets (Supplementary Figure 2).

Models that used SMILES or SMILES-derived features showed a significant difference in their performance ($p$-value of $6.03 \times 10^{-6}$ for RMSE, $p$-value of $3.18 \times 10^{-6}$ for PCC; Kruskal-Wallis test). Among them, *Mol Graph (GCN)* and *SMILES (Transformer)* achieved the lowest median RMSE (14.7 for both) and the highest median PCC (0.87 for both). Both of them performed significantly better than the other models ($p$-value $< 0.02$ for both RMSE and PCC, one-sided Mann-Whitney U test with Benjamini-Hochberg correction). There was no significant difference between these two models ($p$-value of $> 0.25$ for both RMSE and PCC, two-sided Mann-Whitney U test with Benjamini-Hochberg correction).

The models exploiting drug target based features were similar to each other in performance, with median RMSE ranging from 17.63 to 17.93 and PCC of 0.79 ($p$-value $> 0.45$ for RMSE and PCC; Kruskal-Wallis test).

Among the models utilizing gene expression data, *LINCS_1000*, which uses expression from 978 landmark genes, achieved the best performance with a median RMSE of 11.43 and a PCC of 0.95. It significantly outperformed the *Genex* model, which leverages expression data from all available genes (median RMSE of 13.6, PCC of 0.93; adjusted $p$-value = 0.004, one-sided Mann-Whitney U test with Benjamini-Hochberg correction).

Overall, the models demonstrated moderate to strong predictive capabilities for unseen triplets. However, none outperformed the baseline model (adjusted $p$-value of 1.0).

| Dataset Processed by | Model | RMSE (Mean (std)) | PCC (Mean (std)) |
|---|---|---|---|
| MARSY | MARSY | 9.06 (0.45) | 0.84 (0.01) |
| | Baseline | 10.02 (0.23) | 0.81 (0.01) |
| SynergyX | SynergyX | 9.51 (0.27) | 0.91 (0) |
| | Baseline | 10.95 (0.22) | 0.88 (0.005) |

Supplementary Table 1: Performance comparison of the baseline model with MARSY and SynergyX on their respective preprocessed datasets using leave drug pair split.

#### 1.3 Performance of Baseline vs. Two State-of-the-art Models on Model-specific Datasets

We did not evaluate any previously published models directly in our main experiments. However, to gauge the performance of our implemented baseline (using only one-hot encoding as features), we trained and tested it on the preprocessed synergy datasets released by two recent models: MARSY [1] and SynergyX [2]. MARSY is a multitask deep learning model that predicts drug synergy and individual drug responses by combining untreated cell line gene expression profiles with drug-induced differential expression signatures. It employs MLP-based encoders for drug pair and drug–cell line features, which it fuses through a final MLP to produce three outputs: responses for each drug and the synergy score. SynergyX, a transformer-based multimodal model, leverages mutual and self-attention modules to capture drug–drug and drug–cell interactions. It represents drugs using SMILES-derived ESPF fingerprints and incorporates six types of cell line features: gene expression, mutations, copy number, methylation, gene effect, and dependency probability.

We used the preprocessed synergy datasets provided by the authors and split each into training, validation, and test sets using a 3:1:1 ratio using the leave drug pair strategy. We tuned our baseline model on the validation set and reported its performance on the test set using the optimal hyperparameters (Supplementary Table 1). The baseline closely matched the original models' performance: the baseline achieved a PCC of 0.81 as opposed to MARSY's 0.84 and 0.88 versus SynergyX's 0.91 on the corresponding datasets.

Note that we did not rerun MARSY or SynergyX but reported results published by the authors. Although we used the same input data and the same data splitting strategy (i.e., leave drug pair), it is possible that our train–validation–test splits differ from those in the original studies. Nevertheless, the baseline model showed low variation across five independent runs (standard deviations of 0.01 and 0.005 for PCC), indicating that the results remained consistent across different data partitions.

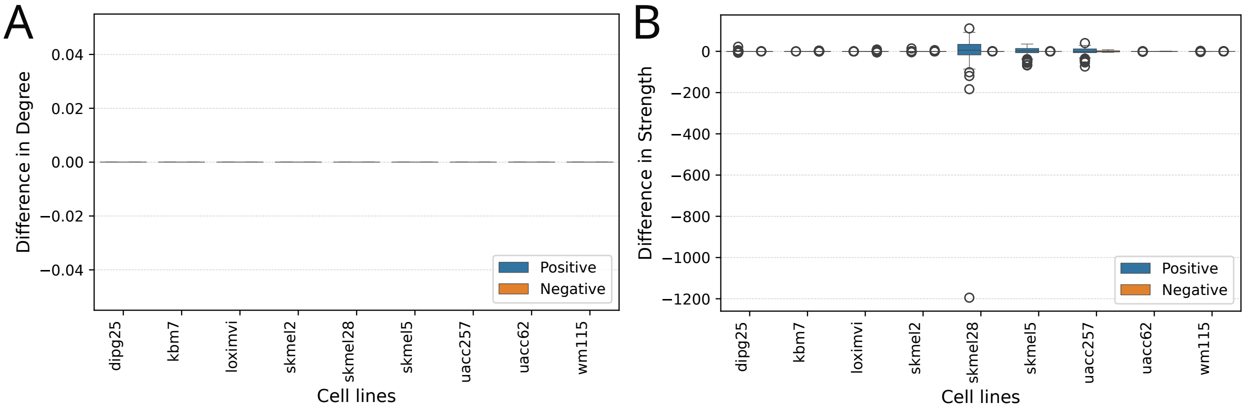


Supplementary Figure 3: **Box plot of differences in degree and strength sequence between original and rewired network. A.** The $x$-axis represents the cell lines. The $y$-axis represents the difference in degree (of each node) between the original and network rewired using the Maslov–Sneppen method. **B.** The $x$-axis represents the cell lines. The $y$-axis represents the difference in strength (of each node) between the original and network rewired using the simulated annealing-based method.

#### 1.4 Network rewiring for network-based ablation study

We used the Maslov-Sneppen [3] randomization method provided in the Brain Connectivity Toolbox (`https://github.com/aestrivex/bctpy`) to rewire the original undirected network while preserving the degree sequence (Supplementary Figure 3A). For strength preserving randomization, we used a simulated annealing-based method [4] (Supplementary Figure 3B). For all these methods, we used approximately 10 swaps per edge.
